# The HPV E7 oncoprotein promotes LIM and SH3 Domain Protein 1 (LASP1) transcription via the Rb/E2F1 signalling pathway in HPV-positive cervical cancer cells

**DOI:** 10.64898/2026.04.08.717142

**Authors:** Miao Wang, Yigen Li, Molly R. Patterson, James A. Scarth, Ethan L. Morgan, Andrew Macdonald

**Affiliations:** School of Molecular and Cellular Biology, Faculty of Biological Sciences, University of Leeds, Leeds, West Yorkshire, LS2 9JT, UK; Astbury Centre for Structural Molecular Biology, University of Leeds, Leeds, West Yorkshire, LS2 9JT, UK; Division of Cancer Therapeutics, The Institute of Cancer Research, London, UK; Genome Regulation and Cell Signaling, Ellen and Ronald Caplan Cancer Center, The Wistar Institute, Philadelphia, Pennsylvania, USA; Barts Cancer Institute, Queen Mary University of London, London, UK; Tumour Virology Group, The Cyprus Institute of Neurology and Genetics, Nicosia, Cyprus

**Keywords:** HPV, cervical cancer, E7, LASP1, E2F1, Rb

## Abstract

Since its discovery in a metastatic lymph node in breast cancer patients, LIM and SH3 Domain Protein (LASP1) has been shown to be over-expressed in and promote the progression of various cancers. We recently demonstrated that LASP1 is highly expressed in human Papillomavirus positive (HPV+) cervical cancers where it promotes cell proliferation and invasion. Importantly, we showed that the HPV E7 oncoprotein increased LASP1 expression by downregulating the microRNA miR-203, which directly targets the *LASP1* mRNA 3’UTR. However, whether LASP1 is regulated by other mechanisms in HPV+ cervical cancers is unclear. Here, we demonstrate an additional mechanism by which HPV E7 regulates *LASP1* transcription. Our data demonstrates an important the role for Rb/E2F1 signalling in promoting LASP1 expression in HPV+ cervical cancer cells. Mechanistically, E7-mediated Rb binding and degradation is required for E7-driven *LASP1* promoter activity. Overexpression of Rb decreased *LASP1* promoter activity, *LASP1* mRNA expression and LASP1 protein levels, whereas E2F1 expression promoted LASP1 expression. Importantly, E2F1 directly bound to the *LASP1* promoter region and the E2F1 binding sites are essential for LASP1 expression in HPV+ cervical cancer cells. Finally, we demonstrate that LASP1 can partially rescue the growth defects observed in E2F1 knockdown cervical cancer cells. Taken together, our data show that HPV E7 regulates LASP1 expression via the Rb/E2F1 signalling pathway. In combination with our previous work, our studies demonstrate that HPV E7 employs multiple mechanisms to drive LASP1 expression, reinforcing the importance of LASP1 in HPV+ cervical cancer.

## Introduction

Persistent infection with high-risk human papillomavirus (HR-HPV) is the principal cause of cervical cancer worldwide and the fourth leading cause of cancer-related deaths in women [1]. The HPV oncoproteins E6 and E7 act synergistically to immortalize and transform infected cells [2–4]. HPV E7 drives S phase progression and cell proliferation via promoting the degradation of the retinoblastoma protein (Rb), releasing the E2F transcription factor [5, 6]. In normal cells, this can activate p53-mediated apoptosis [7]; however, the co-expression of HPV E6, which promotes p53 degradation, blocks apoptosis and thus promotes cell immortalisation [8–10]. The E5 oncoprotein can also contribute to cell proliferation by several mechanisms, including promoting EGFR activity, contributing to the transforming effects of E6 and E7 [11–14]. In addition to these well characterised functions, HPV E6 and E7 target and modulate the activity of multiple host cellular factors to promote proliferation, delay differentiation, suppress apoptosis and evade host immune surveillance [15–19]. Understanding how these oncoproteins interact with host cells may identify new avenues for cervical cancer therapies.

LIM and SH3 Domain-containing Protein (LASP1) is a scaffold protein and focal adhesion adapter protein [20]. Increasing evidence demonstrates a role for LASP1 in the progression of many cancers including breast cancer, colorectal cancer and hepatocellular carcinoma [21–23]. LASP1 contains an amino-terminal LIM domain (named after the initial discovery of the Lin11, Isl-1 and Mec), followed by two nebulin-like repeats (NRs) and a carboxyl-terminal SRC homology 3 (SH3) domain. It is involved in numerous biological and pathological processes, especially in the regulation of dynamic actin-based, cytoskeletal activities and cell motility [20]. The LIM domain contains two zinc finger motifs, suggesting this domain may function to bind DNA and RNA [24]. The SH3 domain can bind to proline rich proteins; this domain has been shown to interact with proline-rich cytoskeletal proteins directly, such as Zyxin and LPP, suggesting LASP1 may have an important role in pseudopodal formation, extension, and invasion [25–27]. In the NR1 region, there is a nuclear export signal (NES; [28]). In the linker region, there are two phosphorylation sites; S146 (serine at position 146) and Y171 (tyrosine at position 171). S146 is phosphorylated by protein kinase A and protein kinase G and is dephosphorylated by protein phosphatase 2B (PP2B), and Y171 is phosphorylated by c-Src and c-Abl non-receptor tyrosine kinases [25, 29, 30]. Functionally, these two sites may be involved in LASP1 re-localization between the cytoplasm and nucleus [28].

Recently we showed that LASP1 is overexpressed in HPV-driven cervical cancer and is critical for tumour growth, both *in vitro* and *in vivo* [31]. We demonstrated that HPV E7 can increase LASP1 expression by suppressing the tumour suppressive microRNA miR-203, which directly targets the LASP1 mRNA. However, whether E7 uses additional mechanisms to regulate LASP1 expression remains unanswered. Here, we report that E7 proteins from high-risk HPV types increase LASP1 transcription through an Rb/E2F1 signalling pathway. We further show that E2F1 controls LASP1 expression by directly binding to the LASP1 promoter, which is the first time E2F1 has been demonstrated to act as a transcriptional regulator of LASP1. These bring new insight into the mechanisms of E7-mediated LASP1 upregulation in HPV-associated cervical cancer.

## Results

### High-risk HPV E7 proteins regulate *LASP1* promoter activity

In our previous study, we demonstrated that HPV E7 could increase LASP1 mRNA expression and protein levels [31]. However, a detailed analysis of E7-mediated LASP1 transcriptional regulation was not performed. To further investigate if HPV E7 regulates LASP1 expression at the transcriptional level, we cloned the *LASP1* promoter sequence, obtained from UCSC Genome Bioinformatics (https://genome.ucsc.edu/), into the luciferase reporter system pGL3.0. The luciferase reporter assay results showed that the transcriptional activity driven by the *LASP1* promoter was significantly enhanced compared to pGL3.0 alone (Figure 1A). Co-expression of E7 proteins from both HR HPV16 and HPV18 increased *LASP1* promoter activity in a dose dependent manner (Figure 1B-D). To investigate if these effects were conserved across HPV types, we tested E7 proteins from both high-risk (HPV16 and 18) and low-risk (LR; HPV6, 8, 11 and 38) types (Figure S1A). Our data show that only HR HPV E7 increases *LASP1* promoter activity (Figure 1D and E), suggesting that the ability to drive *LASP1* transcription may be linked to the oncogenic functions of HPV E7.

**Figure 1.**
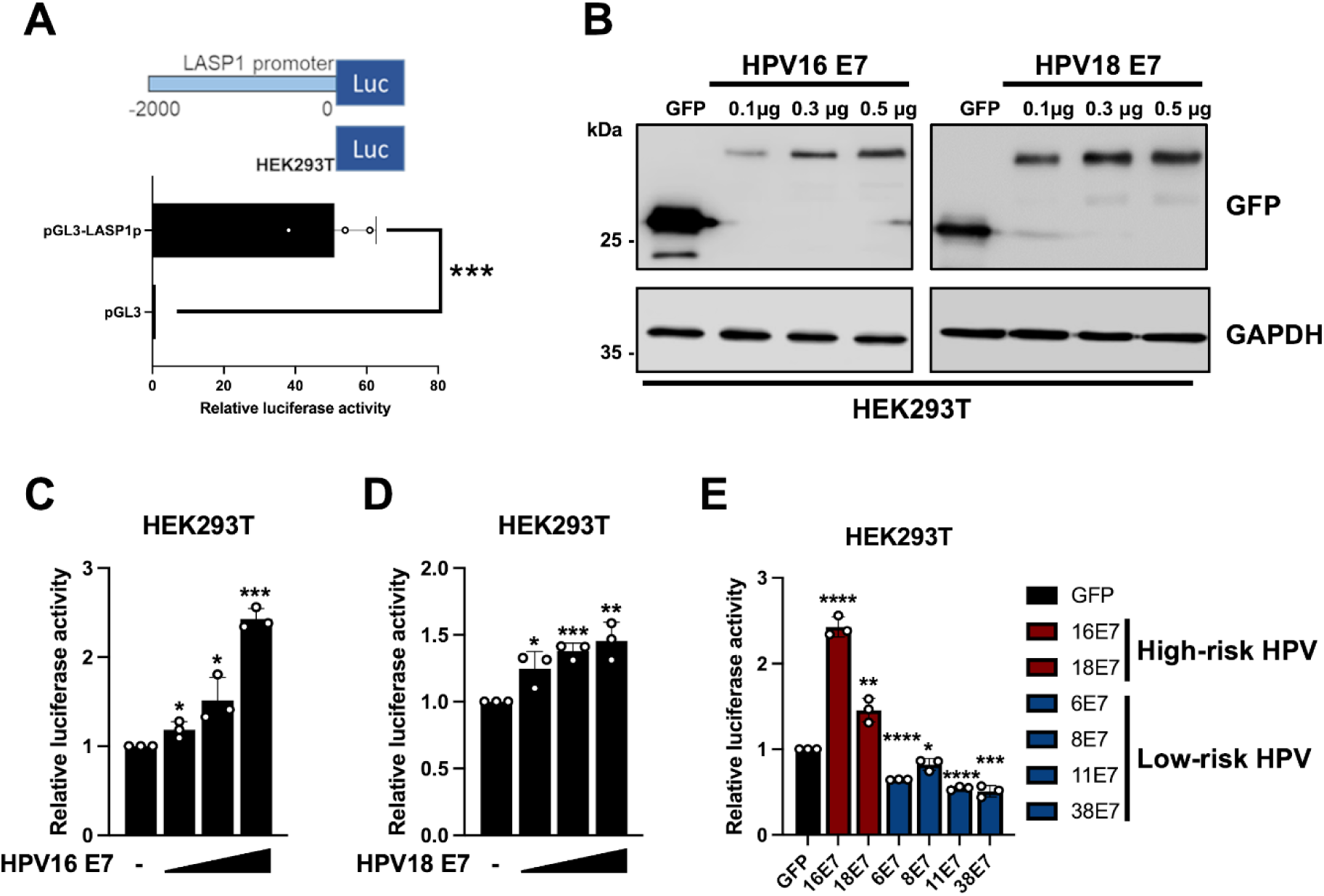
High-risk HPV E7 proteins regulate *LASP1* transcription. (A) Dual luciferase reporter analysis driven by the *LASP1* promoter (pGL3-LASP1p) in HEK293T cells compared to empty vector control (pGL3). Luminescence values of firefly luciferase activity were normalised to *Renilla* luciferase activity. Figure was made using BioRender.com. (B) Representative western blot of HEK293T cells transiently transfected with GFP or increasing amounts of GFP-HPV16 or GFP-HPV18 E7 (0.1 μg, 0.3 μg and 0.5 μg) and analysed for GFP expression. GAPDH served as a loading control. (C-D) Dual luciferase reporter analysis driven by the *LASP1* promoter in HEK293T cells transiently transfected with GFP or increasing amounts of (C) GFP-HPV16 or (D) GFP-HPV18 E7 (0.1 μg, 0.3 μg and 0.5 μg). Luminescence values of firefly luciferase activity were normalised to *Renilla* luciferase activity. (E) Dual luciferase reporter analysis driven by the *LASP1* promoter in HEK293T cells transiently transfected with different types of HPV E7 (0.5 μg GFP-tagged HPV E7). Luminescence values of firefly luciferase activity were normalised to *Renilla* luciferase activity. Data shown are representative of at least three independent experiments. Error bars represent the mean +/− standard deviation of a minimum of three biological repeats. \**P* < 0.05, \*\**P* < 0.01, \*\*\**P* < 0.001, \*\*\*\**P* < 0.0001 (Student’s *t*-test).

### The Rb binding domain of HR-HPV E7 is indispensable for increasing *LASP1* promoter activity

To investigate how HR-HPV E7 increases *LASP*1 promoter activity, a panel of well-characterised E7 mutants were generated by site-directed mutagenesis (Figure 2A). These include HPV16 E7 Δ22-26 (required for Rb and p107 binding [7]; L67R (required for histone deacetylases (HDAC) binding [32]); HPV18 E7Δ24-27 (required for Rb and p107 binding [33]); C27S (required for Rb, but not p107 binding [33]) and SS32/34AA (abrogates CKII mediated phosphorylation [34, 35]). A schematic of HPV16 and 18 E7 proteins is shown in Figure 2A and expression of the mutants is shown in Figure S1B and S1C. When compared to the control, the HPV16 E7 Δ22-26 and HPV18 E7 Δ24-27 or C27S mutants failed to increase *LASP1* promoter activity, while HPV16 E7 L67R and HPV18 E7 SS32/34AA increased activity similar to the wild type (WT) E7 (Figure 2B and C). These results suggest that Rb binding by HR-HPV E7 is required for increased *LASP1* promoter activity.

**Figure 2.**
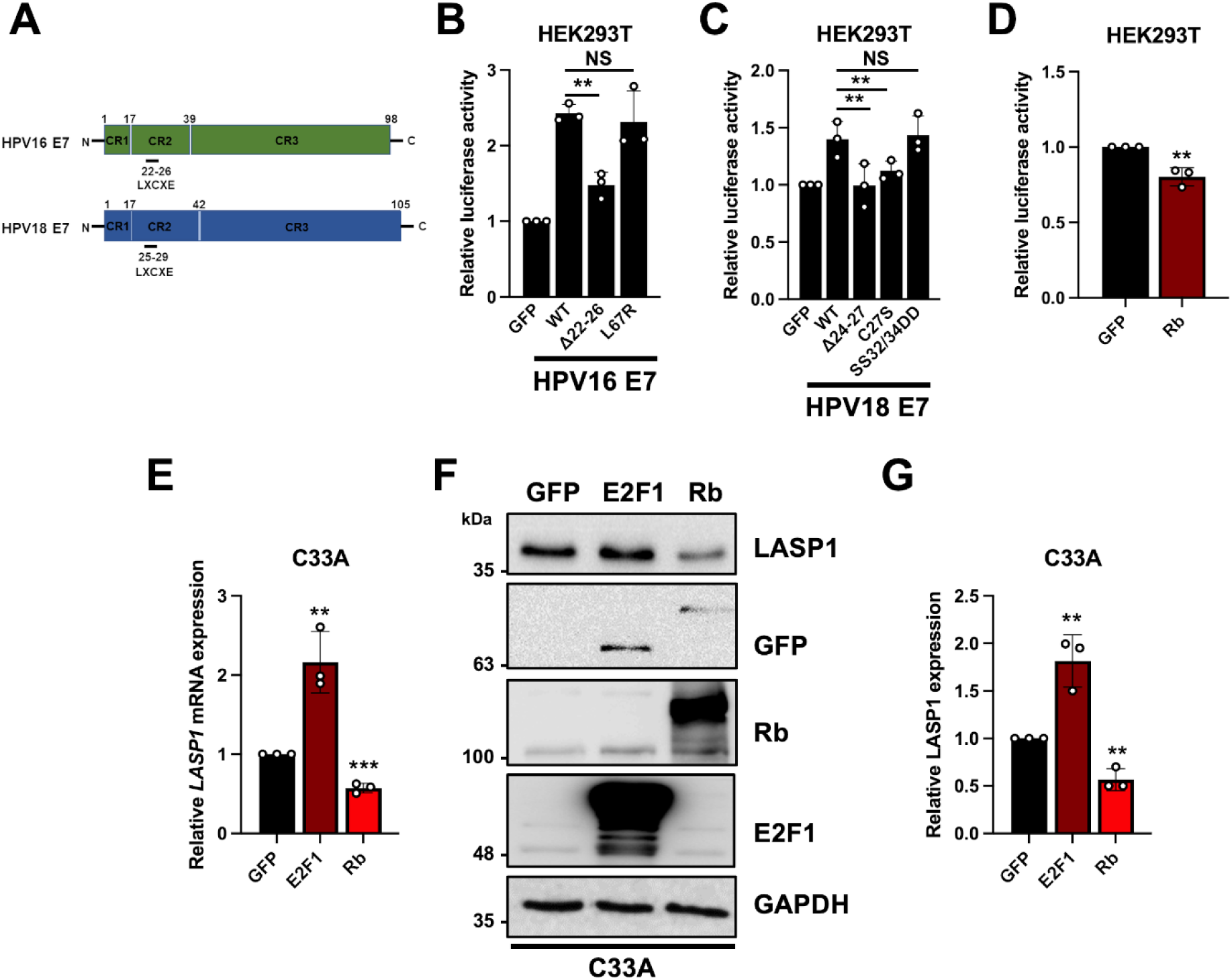
HPV E7 regulates *LASP1* transcription via the RB-E2F1 pathway. (A) Schematic diagram of HPV16 E7 and HPV18 E7 structures. The LXCXE motif is located between amino acids 22-26 in HPV16 E7 and between 25-29 in HPV18 E7. Figure was made using BioRender.com. (B) Dual luciferase reporter analysis driven by the *LASP1* promoter in HEK293T cells transiently transfected with GFP or wild type (WT) HPV16 E7 and the highlighted mutants. Luminescence values of firefly luciferase activity were normalised to *Renilla* luciferase activity. (C) Dual luciferase reporter analysis driven by the *LASP1* promoter in HEK293T cells transiently transfected with GFP or wild type (WT) HPV18 E7 and the highlighted mutants. Luminescence values of firefly luciferase activity were normalised to *Renilla* luciferase activity. (D) Dual luciferase reporter analysis driven by the *LASP1* promoter in HEK293T cells transiently transfected with GFP or Rb. Luminescence values of firefly luciferase activity were normalised to *Renilla* luciferase activity. (E) Relative mRNA levels of *LASP1* in C33A cells transiently transfected with GFP, E2F1 or Rb. *U6* and *GAPDH* served as loading controls. (F-G) Representative western blot of C33A cells transiently transfected with GFP E2F1 or Rb. Lysates were analysed for LASP1, E2F1, Rb and GFP expression. GAPDH served as a loading control. Relative protein expression of LASP1 is shown in G. Data shown are representative of at least three independent experiments. Error bars represent the mean +/− standard deviation of a minimum of three biological repeats. \**P* < 0.05, \*\**P* < 0.01, \*\*\**P* < 0.001 (Student’s *t*-test).

### LASP1 expression is regulated by the Rb/E2F1 pathway

To verify if Rb plays a role in regulating *LASP1* transcription, we overexpressed Rb in cells co-transfected with the *LASP1* promoter reporter plasmid (Figure S1D). Upon co-expression of Rb, *LASP1* promoter activity was significantly reduced (Figure 2D). Furthermore, when overexpressed in HPV-C33A cervical cancer cells, *LASP1* mRNA expression and LASP1 protein levels were significantly reduced (Figure 2E-G). Conversely, overexpression of E2F1, an Rb-binding transcription factor, enhanced LASP1 expression (Figure 2E-G, S1D). These data suggest that the Rb/E2F1 signalling pathway plays a role in the regulation of LASP1 expression.

### HR-HPV E7 promotes LASP1 expression in an E2F1-dependent manner

The transcription factor E2F1 can bind to hundreds of promoters to drive the expression of genes associated with a wide range of cellular pathways [36, 37]. To confirm the role of E2F1 in the regulation of LASP1 expression in HPV+ cancer cell lines, E2F1 was overexpressed in HPV18+ HeLa and HPV16+ SiHa cells. E2F1 overexpression significantly upregulated LASP1 expression at both the mRNA and protein levels (Figure 3A and B). To confirm if HPV E7-mediated LASP1 expression is E2F1-dependent, E2F1 was co-transfected with a pool of HPV E7 siRNAs previously described [31]. As expected, LASP1 expression decreased after the depletion of E7; however, E2F1 overexpression partially rescued LASP1 expression, suggesting that HPV E7-induced LASP1 expression was partially E2F1-dependent (Figure 3C).

**Figure 3.**
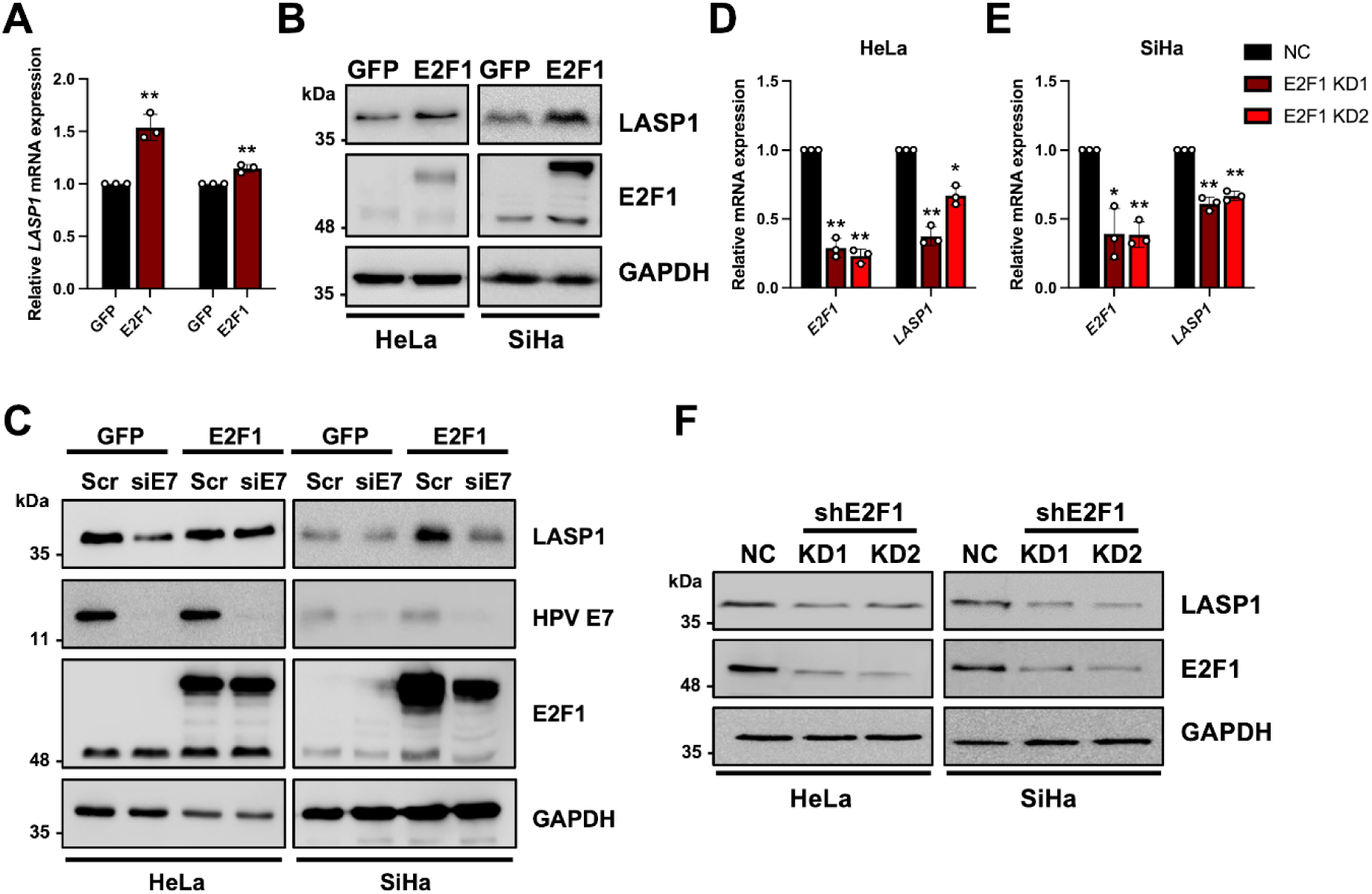
E2F1 promotes LASP1 expression in HPV+ cervical cancer cells. **(A)** RT-qPCR analysis of *LASP1* mRNA level in HeLa and SiHa cells transiently transfected with GFP or GFP-E2F1. *U6* and *GAPDH* served as loading controls. (B) Representative western blot of HeLa and SiHa cells transiently transfected with GFP or GFP-E2F1. Lysates were analysed for LASP1 and GAPDH served as a loading control. (C) Representative western blot of HeLa and SiHa cells transiently transfected with HPV16 E7 or HPV18 E7 specific siRNAs. Cells were also transiently transfected with GFP or GFP-E2F1. Lysates were analysed for LASP1, HPV 16/18 E7 and E2F1. GAPDH served as a loading control. (D-E) RT-qPCR analysis of *LASP1* mRNA level in (D) HeLa and (E) SiHa cells after shRNA mediated depletion of E2F1 (termed E2F1 knockdown (KD)1 and 2). *U6* and *GAPDH* served as loading controls. (F) Representative western blot in HeLa and SiHa E2F1 KD cells. Lysates were analysed for LASP1 and E2F1 expression. GAPDH served as a loading control. NC, negative control. Data shown are representative of at least three independent experiments. Error bars represent the mean +/− standard deviation of a minimum of three biological repeats. \**P* < 0.05, \*\**P* < 0.01, \*\*\**P* < 0.001 (Student’s *t*-test).

To confirm these data, we depleted E2F1 using two different E2F1 short hairpin RNA (shRNA)s in both HeLa and SiHa cells (shRNA sequences shown in Table 1). Depletion of E2F1 in HeLa and SiHa cells was confirmed by RT-qPCR and western blot (Figure 3D-F). Upon E2F1 depletion, LASP1 expression was decreased at both the mRNA and protein level (Figure 3D-F). These data demonstrate that E2F1 regulates LASP1 expression in HPV+ cervical cancer cells.

**Table 1.**
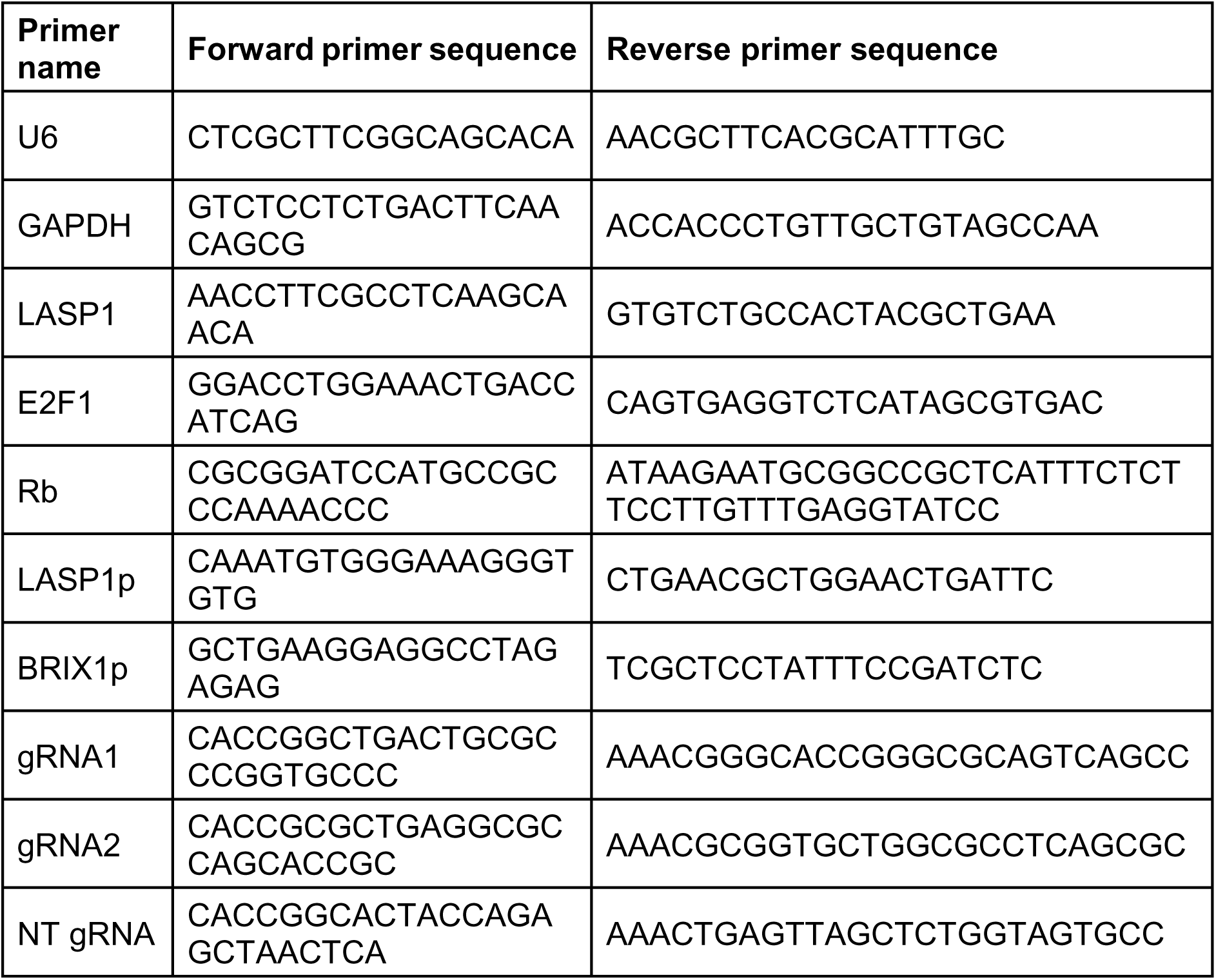
Primers used in this study.

### E2F1 binding to the *LASP1* promoter is required for LASP1 expression in HPV+ cervical cancer cells

To further verify the role of E2F1 in regulating the LASP1 expression, E2F1 and the *LASP1* promoter reporter plasmid were co-transfected into HEK293T cells. Upon increasing E2F1 expression, *LASP1* promoter activity was enhanced in a dose-dependent manner (Figure 4A and B). Consistent with this, a significant decrease in *LASP1* promoter activity was observed in E2F1 KD HeLa and SiHa cells, indicating that E2F1 drives *LASP1* promoter activity in HPV+ cervical cancer cells (Figure 4C and D).

**Figure 4.**
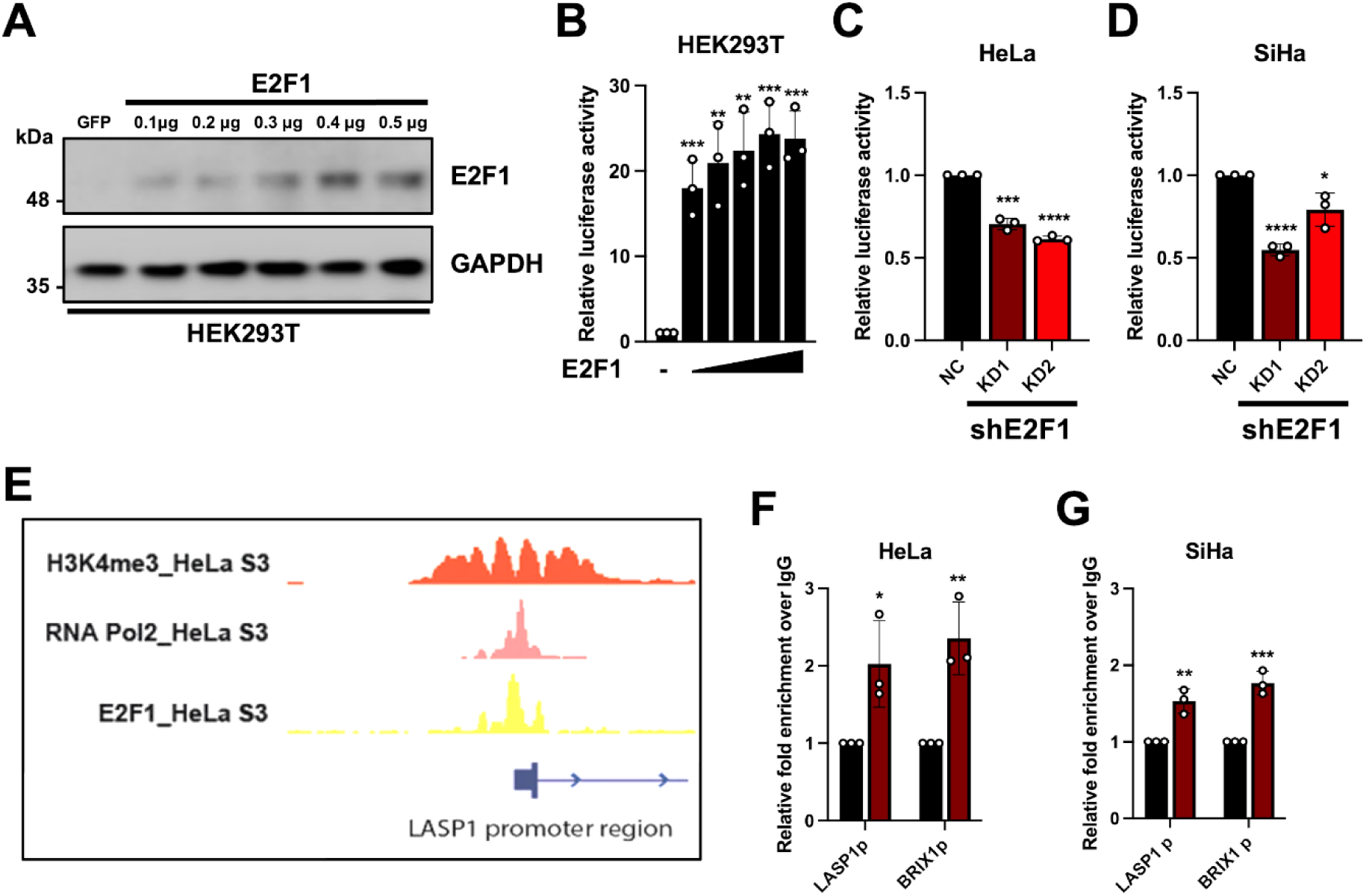
E2F1 binds to the *LASP1* promoter. (A) Representative western blot of HEK293T cells transiently transfected with GFP or GFP-E2F1. Lysates were analysed for E2F1 and GAPDH served as a loading control. (B) Dual luciferase reporter analysis driven by the *LASP1* promoter in HEK293T cells transiently transfected with increasing amounts of E2F1 (0.1 μg, 0.2 μg, 0.3 μg, 0.4 μg and 0.5 μg). Luminescence values of firefly luciferase activity were normalised to *Renilla* luciferase activity. (C-D) Dual luciferase reporter analysis driven by the *LASP1* promoter in (C) HeLa and (D) SiHa E2F1 KD cells. (E) ChIP-seq data of H3K4me3, RNA polymerase II and E2F1 from HeLa S3 cells, mapped to the promoter region of *LASP1*. Data was visualised using Integrative Genomics Viewer. (F-G) ChIP-qPCR analysis of E2F1 binding at the *LASP1* promoter in (F) HeLa and (G) SiHa cells. All values at LASP1p and BRIX1p (positive control region) were normalised to the input. Fold enrichment relative to IgG bound samples is shown. NC, negative control. Data shown are representative of at least three independent experiments. Error bars represent the mean +/− standard deviation of a minimum of three biological repeats. \**P* < 0.05, \*\**P* < 0.01, \*\*\**P* < 0.001 (Student’s *t*-test).

To identify potential E2F1 binding sites in the *LASP1* promoter region, we performed bioinformatic analysis. The published ChIP-seq data of E2F1, RNA polymerase II and histone modifications (H3K4me3) from HeLa S3 cells, a subclone derived from the parental HeLa cell line, were mapped onto the *LASP1* promoter region and visualized using Integrative Genomics Viewer (Figure 4E). The source of the ChIP-seq data shown in Figure 4E are listed in Table 2. H3K4me3 is highly enriched at active promoters [38]. Both RNA polymerase II and H3K4me3 were found enriched at the *LASP1* promoter region (2kb upstream of start codon) suggesting an active promoter; interestingly, E2F1 was also enriched at the *LASP1* promoter, suggesting that it may be directly involved in *LASP1* transcription. To confirm this, we performed ChIP-qPCR analysis in HeLa and SiHa cells. The *BRIX1* promoter was previously shown to be enriched with E2F1 and was used as a positive control (23). In line with the ChIP-seq data, E2F1 was enriched at the *LASP1* promoter in both HeLa and SiHa cells (Figure 4F-G). These data together suggest that E2F1 directly binds the *LASP1* promoter.

**Table 2.** Accession numbers of GSE dataset used in this study.

| Accession number | Data | Tissue Type |
| --- | --- | --- |
| GSM935484 | E2F1 ChIP-seq data | HeLa S3 cells |
| GSM935395 | RNA Pol II ChIP-seq data | HeLa S3 cells |
| GSE20303 | H3K4me3 ChIP-seq data | HeLa cells |

We next identified the E2F1 binding sites in the *LASP1* promoter using LASAGNA-Search 2.0 in combination with the published ChIP-seq data presented in the Table 2 [39]. A putative E2F1 binding sequence was located at CCTGAGAGCGCTGAG. To investigate the importance of this possible E2F1 binding site, we generated a deletion mutant in the Luciferase reporter construct lacking this sequence using site-directed mutagenesis (Figure 5A). When compared to the WT promoter, the E2F1 binding site mutant had significantly reduced promoter activity (Figure 5B). Moreover, E2F1 only partially increased luciferase expression driven by the mutated promoter compared to the wild type of the *LASP1* promoter (Figure 5B), indicating that E2F1 drives *LASP1* promoter activity via the E2F1 binding sites. To validate these findings, we deleted the E2F1 binding sites in the endogenous *LASP1* promoter in HeLa and SiHa cell lines using CRISPR/Cas9 (Figure 5C). Deletion of the E2F1 binding sites in the *LASP1* promoter resulted in deceased expression of LASP1 at both the mRNA and protein (Figure 5D-F). These data demonstrate that binding of E2F1 to the *LASP1* promoter at the identified E2F1 binding site is required to drive *LASP1* promoter activity and subsequent expression in HPV+ cervical cancer cells.

**Figure 5.**
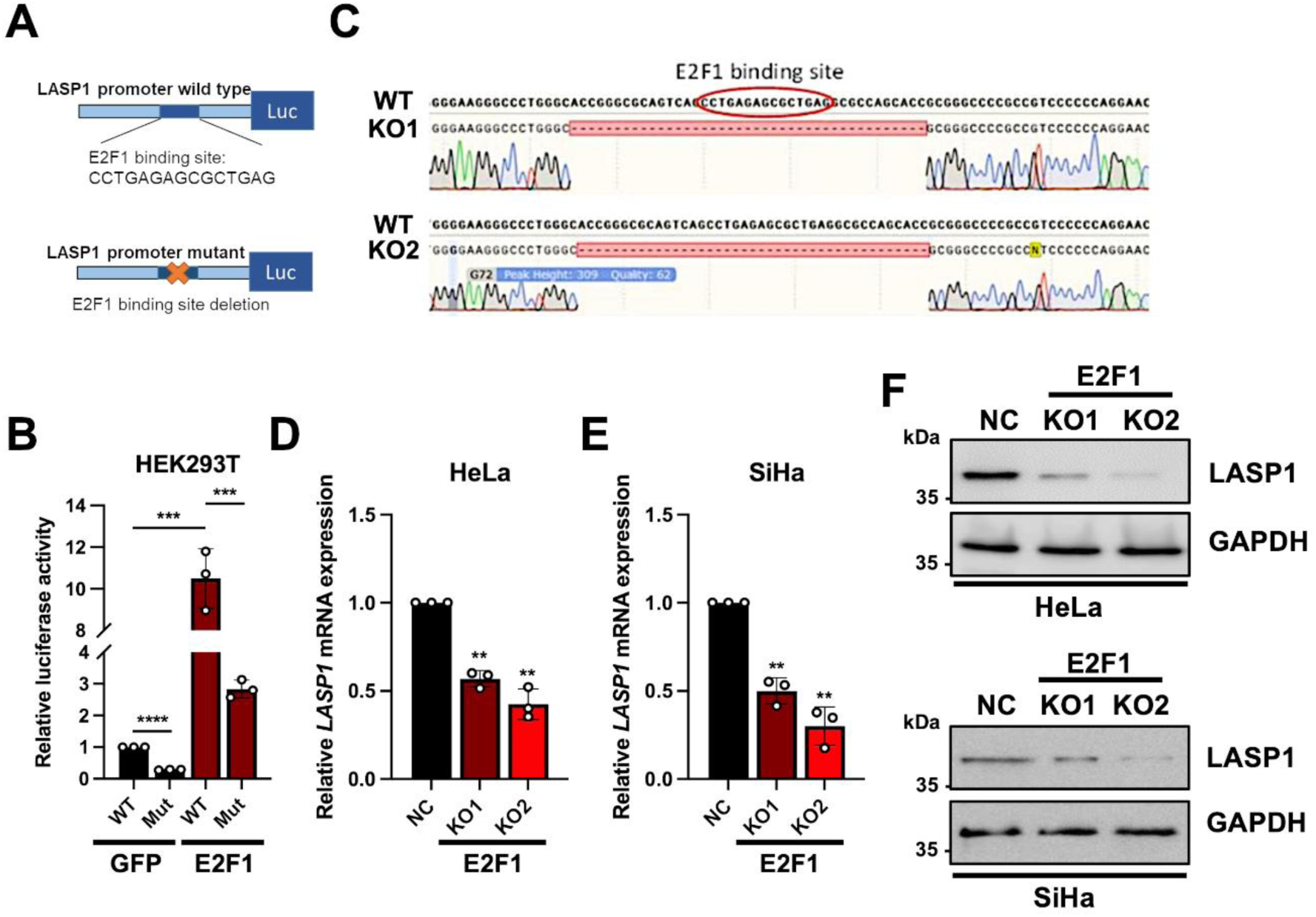
E2F1 binding sites in the *LASP1* promoter are required LASP1 expression in HPV+ cervical cancer cells. (A) Schematic diagram of E2F1 binding motif in the *LASP1* promoter driven luciferase reporter. Figure was made using BioRender.com. (B) Dual luciferase reporter analysis driven by the WT or E2F1 mutant *LASP1* promoter in HEK293T cells transiently transfected with GFP or GFP-E2F1. Luminescence values of firefly luciferase activity were normalised to *Renilla* luciferase activity. (C) Sequencing analysis of HeLa cells after CRISPR-Cas9 mediated knockout (KO) of the E2F1 binding sites in the *LASP1* promoter (E2F1 (termed E2F1 KO1). (D-E) RT-qPCR analysis of *LASP1* mRNA level in (D) HeLa and (E) SiHa cell E2F1 KO cells. *U6* and *GAPDH* served as loading controls. (F) Representative western blot of HeLa and SiHa E2F1 KO cells. Lysates were analysed for LASP1 and GAPDH served as a loading control. NC, negative control. Data shown are representative of at least three independent experiments. Error bars represent the mean +/− standard deviation of a minimum of three biological repeats. \**P* < 0.05, \*\**P* < 0.01, \*\*\**P* < 0.001 (Student’s *t*-test).

### Restoration of LASP1 partially rescues proliferation defects in E2F1 depleted HPV+ cervical cancer cells

The HR-HPV E7 oncoproteins promote cancer cell proliferation, at least in part by up-regulating E2F1-dependent transcription [40]. To determine if LASP1 expression can rescue the proliferation defect in E2F1 KD cells, we overexpressed LASP1 in E2F1 KD HeLa and SiHa cells (Figure 6A-B). Reduction in E2F1 expression caused by shRNA mediated knockdown significantly decreased cell growth and colony formation in HPV+ cervical cancer cells. LASP1 overexpression partially rescued this growth defect in both cell lines (Figure 6C-H). These results reveal that LASP1 plays a role in E2F1-mediated cell proliferation in HPV+ cervical cancer cells.

**Figure 6.**
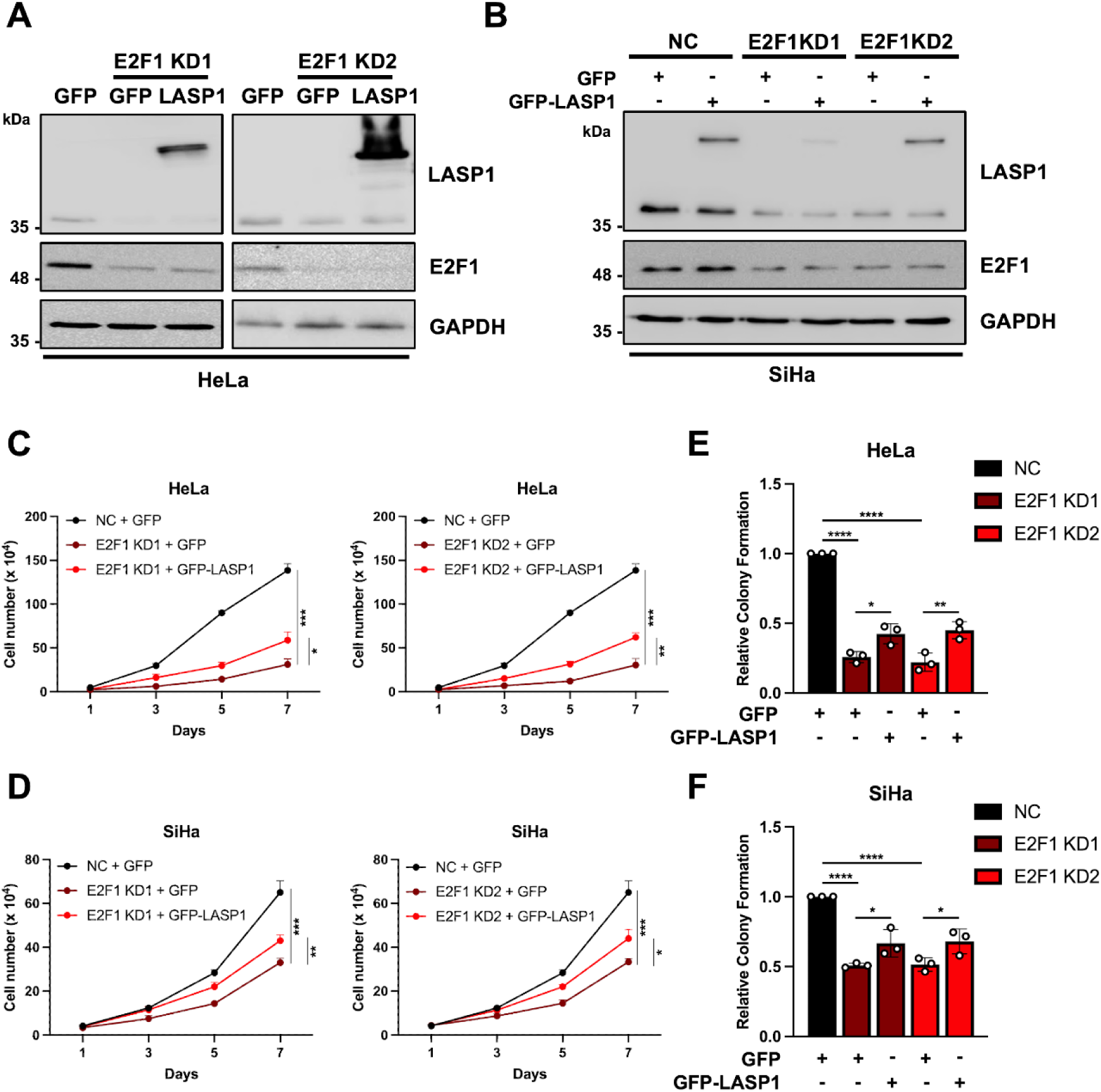
Restoration of LASP1 partially rescues proliferation defects in E2F1 depleted HPV+ cervical cancer cells. (A-B) Representative western blot of (A) HeLa and (B) SiHa E2F1 KD cells transiently transfected with GFP or GFP-LASP1. Lysates were analysed for LASP1 and E2F1 expresion. GAPDH served as a loading control. (C-D) Growth curve analysis of (C) HeLa and (D) SiHa E2F1 KD cells transiently transfected with GFP or GFP-LASP1. (E-F) Colony formation assays of (E) HeLa and (F) SiHa E2F1 KD cells transiently transfected with GFP or GFP-LASP1. NC, negative control. Data shown are representative of at least three independent experiments. Error bars represent the mean +/− standard deviation of a minimum of three biological repeats. \**P* < 0.05, \*\**P* < 0.01, \*\*\**P* < 0.001 (Student’s *t*-test).

## Discussion

In this study, we identified a novel mechanism by which HR-HPV E7 increases LASP1 expression. Our findings build on our previous work, where we established that the cytoskeletal protein LASP1 is overexpressed in HPV-positive cervical cancer, driving tumorigenesis *in vitro* and *in vivo* [31]. Additionally, we demonstrated that HPV E7 upregulates LASP1 by repressing miR-203, which directly targets the *LASP1* 3’ UTR [31].

Here, we further elucidate how HR-HPV E7 drives LASP1 expression at the transcriptional level through Rb degradation and subsequent activation of E2F1, a transcription factor known to regulate cell cycle progression [41]. Interestingly, our results indicate that while HR-HPV E7 proteins activate the *LASP1* promoter, LR-HPV E7 proteins exhibit a small, but significant suppressive effect. This difference may stem from variations in Rb binding affinity between HR- and LR-HPV E7 proteins; HR-HPV E7 binds Rb more tightly, leading to more efficient Rb degradation and enhanced E2F1 activation [42]. Conversely, the weaker Rb binding affinity of LR-HPV E7 could allow for interactions with other pocket proteins, such as p130, potentially recruiting repressive E2F factors like E2F4 and E2F6 to the promoter region [43]. Future studies are required to understand the potential role of LR-HPV E7 proteins in modulating LASP1 expression.

Additionally, we demonstrate that the Rb-binding domain of HR-HPV E7 is essential for upregulating *LASP1* promoter activity. By employing site-directed mutagenesis to generate specific HPV16 and HPV18 E7 mutants, we observed distinct effects on *LASP1* promoter activity. Notably, the Rb-binding-deficient mutants, HPV16 E7 Δ22-26 and HPV18 E7 Δ24-27 and C27S, failed to enhance *LASP1* promoter activity, indicating that Rb binding is indispensable for this effect. In contrast, the mutants HPV16 E7 L67R and HPV18 E7 SS32/34AA, which retain Rb binding but affect other interactions, including HDAC and CKII, were still able to activate the *LASP1* promoter similarly to the wild-type E7 proteins. This suggests that, while other domains of E7 may contribute to cellular regulation, the interaction with Rb is a primary requirement for *LASP1* promoter activation. These findings highlight a novel role of E7’s Rb-binding function in modulating LASP1 expression, contributing to HPV-driven oncogenesis by promoting cell proliferation and metastasis.

The transcriptional regulation of LASP1 is complex, with several transcription factors including HIF1-α, SOX9 and c-Jun, and microRNAs, including miR-145, miR-203 and miR-218, regulating *LASP1* transcription in different tumour types [26, 27, 44–47]. However, some studies produced conflicting results. The tumour suppressor p53 suppressed *LASP1* transcription in hepatocellular carcinoma cells, with the oncogenic p53 mutant R175H abolishing this repression [21]. However, p53 mutations were not associated with LASP1 expression in breast cancers [48]. Thus, the transcriptional regulation of *LASP1* is complex and may be dependent on the genetic background of the cancer. Since the Rb binding site mutants of HR-HPV E7 showed decreased luciferase activity driven by the *LASP1* promoter, we hypothesised that E2F transcription factors may be involved in Rb-mediated *LASP1* transcription. Indeed, published ChIP-seq data from HeLa S3 cells found enrichment of E2F1 in the *LASP1* promoter region (Figure 4E). These data support the mechanism that the E7-mediated LASP1 upregulation in HPV+ cancer cells is at least partially E2F1-dependent. ChIP-qPCR analysis of E2F1 confirmed binding to the *LASP1* promoter and knock-out of the E2F1 binding sites by CRISPR-Cas9 technique significantly reduced *LASP1* transcription. However, E2F1 binding site knock-out did not abolish *LASP1* transcription, suggesting that other transcription factors may be involved that require further exploration.

LASP1 has been documented to play a vital role in regulating cell proliferation in HPV-related cervical cancer cells, as well as in other cancer types [31, 44, 49, 50]. We found that LASP1 can partially rescue cell proliferation even in the absence of E2F1, which is known to regulate the cell cycle progression [41]. Together with our results showing that E2F1 controls *LASP1* transcription, this suggests that LASP1 is regulated by E2F1 to support cellular proliferation. This may be due to the activation of other cellular pathway to drive cell cycle progression and proliferation, which LASP1 has been shown to activate in other cancers [50–52]. Additionally, LASP1’s role in regulating the cytoskeleton could contribute to this rescue by influencing cytoskeletal dynamics [53], a process that can modulate gene expression (such as epithelial and mesenchymal transition related genes) and cell cycle progression. Future studies aimed at dissecting which signalling pathways activated by LASP1 enhance cell proliferation driven by E2F1 in cervical cancer are therefore warranted.

In summary, our data demonstrate an additional layer of regulation of LASP1 by HPV E7 in cervical cancer cells. We show that HR-HPV E7 drives *LASP1* promoter activity and transcription in a manner dependent on Rb degradation and the transcription factor E2F1, which directly binds to *LASP1* promoter (Figure 7). Combined with our previous work, these studies demonstrate the important role of LASP1 in HPV+ cervical cancer and a deeper understanding of LASP1-mediated oncogenic mechanisms in these cancers may uncover therapeutic vulnerabilities to be exploited.

**Figure 7.**
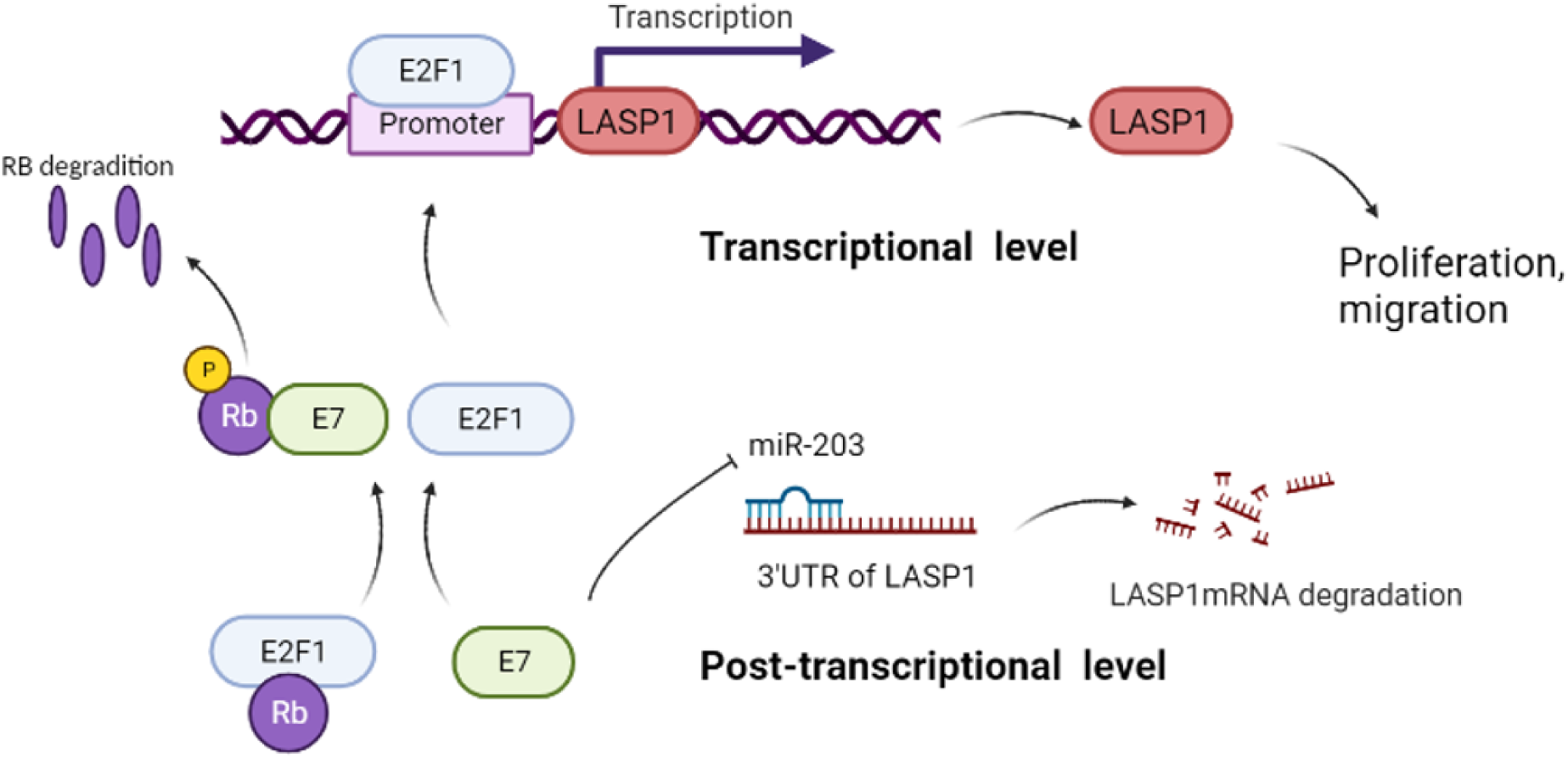
Schematic diagram of HPV E7 mediated activation of LASP1 in HPV+ cervical cancer cells. HPV E7 regulates LASP1 expression via Rb-E2F1 signalling pathway (at the transcriptional level) or by inhibiting miR-203 (at the post-transcriptional level). Figure was made using BioRender.com.

## Materials and Methods

### Cell culture

C33A (HPV negative cervical squamous carcinoma cells) (HTB-31), SiHa (HPV16+ cervical squamous carcinoma cells) (HTB-35), HeLa (HPV18+ cervical epithelial adenocarcinoma) (CRM-CCL-2) and HEK293T cells (CRL-3216) were purchased from ATCC and cultured in Dulbecco’s Modified Eagle’s media (DMEM, Sigma) supplemented with 10 % foetal bovine serum (ThermoFischer Scientific), 50 U/ml Penicillin and 50 μg/ml Streptomycin (Lonza) in a humidified 37°C incubator with 5% CO_2_ and were performed with routine mycoplasma test.

Stable E2F1 knockdown cells using shRNAs were generated as previously described [54]. Briefly, pPAX2, pVSVG and either a pLKO.1-non-targeting shRNA construct or targeting shRNA constructs (sequences available in Table 1) were transfected into HEK293T cells at a ratio of 0.65:0.65:1.2 respectively. 72 h post transfection, complete DMEM media containing lentivirus was removed, filtered and added to the appropriate cell line in a 1:1 ratio with fresh complete DMEM media to cells along with 4 µg/µl Polybrene (Santa Cruz, sc-134220) and 20 mM HEPES. Cells were incubated for 72 h before the addition of selection antibiotic puromycin. Stable cell lines were passaged for 1 week before being screened by RT-qPCR or western blot for successful E2F1 knockdown.

### Plasmids, siRNA and reagents

Plasmids for different HPV E7s are constructed tagged with GFP cloned into pcDNA3.1 using KpnI and EcoRI [55]. Pool of 4 siRNAs targeting LASP1 were obtained from QIAGEN and the siRNAs targeting HPV16 E7 and HPV18 E7 were previously described [31]. The promoter of *LASP1* was cloned into pGL3.0 using NheI and HindIII. Mutagenesis was performed by PCR using the Site-directed mutagenesis kit (NEB). eGFP-tagged LASP1 has been previously described [31] and E2F1 and Rb were respectively cloned into pcDNA3.1 using BamHI and NotI. Lipofectamine^TM^ 2000 (11668019, Invitrogen) or X-tremeGENE™ (6366236001, Roche) were used for transient transfections.

### Cell proliferation assay

Cell growth curves and colony formation assay were previously described [56]. Cells were trypsinised and re-seeded and were counted manually using a haemocytometer. Colonies were stained with crystal violet staining solution (1% crystal violet, 25% methanol) and counted manually.

### Western blot analysis

Separated proteins on the SDS-PAGE gels were transferred to the GE Healthcare Life Sciences™ Hybond nitrocellulose membranes using a Bio-Rad Trans-blot TuRbo transfer system (Bio-Rad), followed by immunoblot analysis. Antibodies used for western blot assay were as follows: HPV16 E7 (sc-65711, Santa Cruz), HPV18 E6 (sc-365089, Santa Cruz), HPV18 E7 (ab100953, Abcam), GAPDH (G-9, SCBT), GFP (sc-9996, SCBT), LASP1 (sc-374059, SCBT), E2F1 (sc-137059, SCBT), Rb (sc-74562, SCBT), HRP conjugated secondary anti-mouse antibodies (Stratech 115-035-174-JIR) and HRP conjugated secondary anti-rabbit antibodies (Stratech 211-032-171-JIR). Colour development with ECL solution (Thermo/Pierce) was performed and the blots were imaged using G:BOX ChemiXT4 system (Syngene).

### RNA extraction and RT-qPCR

Cultured cells were collected using cell scrapers or by trypsin digestion. After centrifugation, total RNA was extracted from cell pellets using an E.Z.N.A total RNA kit I (Omega Bio-tek) according to the manufacturer’s protocol as previous [57]. RNA concentration was measured by a Nanodrop One spectrophotometer (ThermoFisher cientific). After DNase I treatment (ThermoFisher Scientific), mRNA expression levels were detected using GoTaq 1-Step RT-qPCR system (Promega) on a CFX Connect Real-Time PCR detection system (BioRad). Primer sequences can be found in Table 2. The ΔΔCT method was used to analyse relative mRNA expression normalised to the housekeeping genes, *U6* and *GAPDH*. Specific primers were used for each gene analysed and are shown in **Table 1**.

### Dual-luciferase reporter assays

Cells were seeded in 12-well plates and transfected using X-tremeGENE. The luciferase assay using the Dual-Luciferase Kit (Progema) was carried out according to manufacturer’s instructions as previous [58]. Cells were harvested and washed with cold PBS and then resuspended in 150 µl Passive Lysis Buffer. After 15-minute gentle shaking at room temperature, 10 µl of the lysate was added into a luminometer plate and 50 µl of Luciferase Assay Reagent II was added, followed by determination of firefly luciferase activity using a luminometer (EG&G Berthold). Determination of the Renilla luciferase activity as a control was achieved by addition of 50 µl of Stop & Glo reagent. The relative luciferase activity was calculated by normalized to that of mock treated cells.

### Chromatin immunoprecipitation (ChIP) assays

Cells were seeded in 10-cm dishes and harvested at a density of 8×10^6^ cells, followed by the ChIP assay previously described [19]. Briefly, crosslinking was achieved by adding formaldehyde at a final concentration of 0.8 % with gentle 10-minute agitation and was quenched by adding glycine (final concentration is 0.125 M) and with 5-minute agitation at room temperature. The chromatin was sheared into 400∼600 bp fragments using a sonicator at 40% amplitude for eight cycles of 15 seconds with 1 minute ice incubation between each burst. Chromatin samples were purified using thrice phenol/chloroform extraction and were analysed by qPCR.

## Statistical Analysis

Statistical analysis was analysed using GraphPad Prism 8.0 with two-tailed unpaired Student’s T-test. p-value * < 0.05, ** < 0.01, *** < 0.001 and **** < 0.0001. Image J software was used for western blotting densitometry analysis. Unless stated all data presented are representative of at least 3 biological repeats.

## Author Contributions

Conceptualization, MW and AM; Formal analysis, MW; Investigation, MW; Methodology and resources, MW, YL, MRP, JAS, ELM and AM; Writing - original draft, MW; Writing – review & editing, MW, YL, ELM and AM; Supervision, ELM, AM; Project administration, AM; Funding acquisition, AM.

---

**Supplementary Figure 1.**
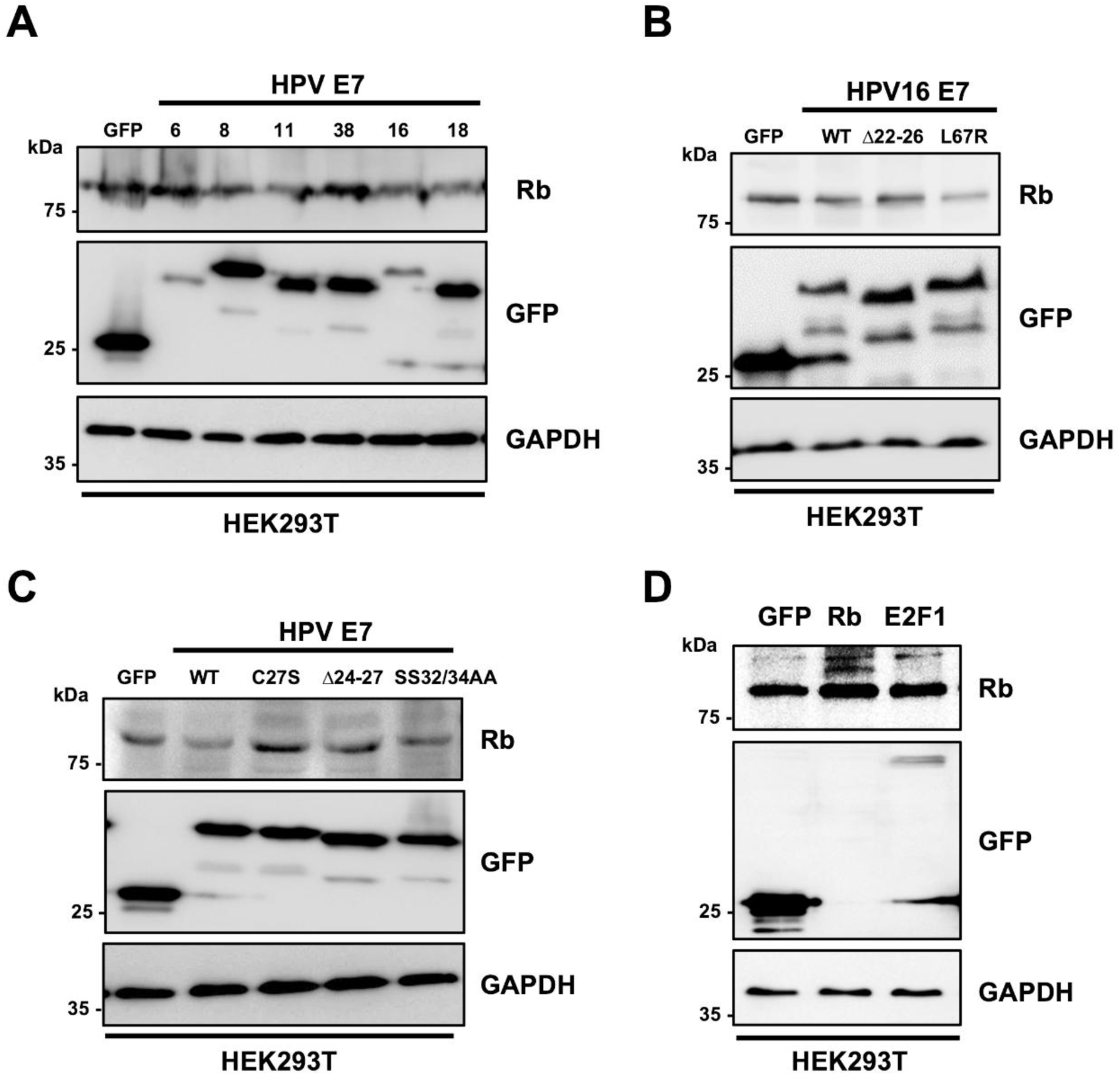
Expression of HPV E7 proteins in HEK293T cells. (A) Representative western blot of HEK293T cells transiently transfected with different types of HPV E7 (0.5 μg GFP-tagged HPV E7). Lysates were analysed for GFP and Rb expression. GAPDH served as a loading control. (B) Representative western blot of HEK293T cells transiently transfected with GFP or wild type (WT) HPV16 E7 and the highlighted mutants. Lysates were analysed for GFP and Rb expression. GAPDH served as a loading control. (C) Representative western blot of HEK293T cells transiently transfected with GFP or wild type (WT) HPV18 E7 and the highlighted mutants. Lysates were analysed for GFP and Rb expression. GAPDH served as a loading control. (D) Representative western blot of HEK293T cells transiently transfected with GFP or Rb. Lysates were analysed for GFP and Rb expression. GAPDH served as a loading control. Data shown are representative of at least three independent experiments.

